# Optimizing sampling for surface localization in 3D-scanning microscopy

**DOI:** 10.1101/2022.03.21.485110

**Authors:** Marie-Anne Burcklen, Frédéric Galland, Loïc Le Goff

## Abstract

3D-scanning fluorescence imaging of living tissue is in demand for less phototoxic acquisition process. For the imaging of biological surfaces, adaptive and sparse scanning schemes have been proven to efficiently reduce the light dose by concentrating acquisitions around the surface. In this paper, we focus on optimizing the scanning scheme at constant photon budget, when the problem is to estimate the position of a biological surface whose intensity profile is modeled as a Gaussian shape. We propose an approach based on the Cramér-Rao Bound to optimize the positions and number of scanning points, assuming signal-dependant Gaussian noise. We show that the optimization problem can be reduced to a few parameters, allowing to define quasi-optimal acquisition strategies, first when no prior knowledge on the surface location is available, and then when the user has a prior on this location.

## 1. INTRODUCTION

State of the art techniques for 3D-scanning fluorescence microscopy provide high-resolution volumetric images of living tissues. One current limitation lies in the fact that the imaging process is phototoxic and causes photobleaching, as labeled tissues are highly irradiated by excitation light [1]. To reduce photodamage, one solution is to engineer the light dose, either by using pulsed light to avoid the deleterious triplet-state of fluorophores [2], or by using a fast feedback to send just enough light to reach a prescribed signal to noise ratio [3–7]. One can also lower the intensity of excitation light, and couple this reduction with denoising to compensate for the reduced signal [1, 8, 9]. Another solution is to change the setup architecture, such as with the light-sheet microscope that allows to excite only the imaged plane [10]. Eventually, instead of performing a standard plane-by-plane acquisition, one can follow an alternative acquisition scheme, such as axial compressed-sensing [11].

In all of these techniques, the scanning process samples the entire 3D volume. To further reduce the light dose, an idea is to shine light only on points that would provide information of interest. In this regard, an adaptive-scanning fluorescence microscope optimized for the unsupervised imaging of biological surfaces has been designed in [12]. It consists in i) performing a fractional pre-scan of the volume in order to estimate the surface of interest, and then ii) targeting illumination inside a thin shell enclosing this surface. In the case of epithelia – curved cell monolayers –, the pre-scan is performed on a very small fraction (~ 0.1 %) of the sample space, and therefore contributes very little to the total light dose. Focusing the scan-path exclusively inside the shell allows to dramatically improve the photon budget. A similar strategy is adopted in [13] to estimate the surface of a cell sheet from the two-photon fluorescence emitted in Lissajou pre-scans.

In the aforementioned work, the axial position of the surface is estimated from the set of pre-scanning points. In light of the above, the quality of this estimation is crucial. Indeed, an accurate estimation allows to restrict the shell as close as possible to the actual structure of interest. But, meanwhile, this accurate estimation must be done with as few photons as possible.

The estimation accuracy depends strongly on the sampling strategy adopted during the prescan. In this paper, we search for the sampling scheme that provide the best trade-off between accuracy and light dose. The estimation problem is the following: for a selected couple of lateral coordinate (*x, y*), our goal is to determine the position *z_s_* of the biological surface along the z-axis. We assume that we have a fixed photon budget to distribute along *z*. The question we address is then: for a given (*x, y*), what are the positions and the number of scanning points along *z* that provide the most accurate estimation of *z_s_*, for a given photon budget? Is it preferable to concentrate this budget on few sampling points along *z*, with relatively high intensity on each scanning point, or to perform many acquisitions along *z* with very few photons per acquisition?

Nevertheless, the answer is not trivial, since the problem depends on many parameters, such as the intensity of the fluorescent signal coming from the tissue, the presence of a background signal, the kind of noise and its level in the imaging system, but also on the fact that the user can have – or not – prior knowledge on the surface’s location. In this paper, we propose to optimize the sampling strategy using the Cramér-Rao Bound (CRB) that is commonly used to characterize estimation performance [14] and to optimize imaging systems [15–17]. The CRB is a lower bound of the variance of estimation for unbiased estimation algorithms. In our case, the CRB is used to assess for the localization accuracy. We show that, when the fluorescent signal along *z* is modeled as a Gaussian shape – which constitutes an accurate approximation in many cases –, we can design optimized sampling strategies that only depend on few reduced parameters. We first study the situation where we have no prior knowledge on the position of the biological surface. We then cover the case where we have a prior knowledge of its approximate position. Such a prior may stem from an iterative procedure to delineate the surface – the accuracy increasing with each iteration –, or simply from the previous time point in a live imaging context.

The paper is organized as follows. Section 2 describes the model of the noisy fluorescent signal along *z*, and the general expression of the CRB of the surface’s position at fixed photon budget. In Section 3, we analyze the CRB when assuming no prior knowledge on the surface’s location, while section 4 tackles the case knowing approximately the surface’s location.

## 2. MODEL AND BOUND ON THE ESTIMATION OF THE AXIAL POSITION OF A BIOLOGICAL SURFACE

### A. Model of the fluorescent signal

In this study, we consider a biological surface labeled with fluorescent tags. Typical biological surfaces are epithelium that are layered cell sheets. An example of such an epithelium is shown in Fig 1.a. It consists of a wing imaginal disk of Drosophila melanogaster. The imaginal disk is the precursor, inside the developing larva, of the wing and thorax of the adult fly. It is an important model system to study the regulation of growth and morphogenesis [18]. In Fig. 1.a, the E-cadherin protein involved in the adherens junction between cells is tagged with the green fluorescent protein (GFP) [19]. It is imaged using a conventional confocal spinning-disk microscope coupled with an EM-CCD sensor. A lateral section of the volumetric image is given in Fig. 1.b.

**Fig. 1.**
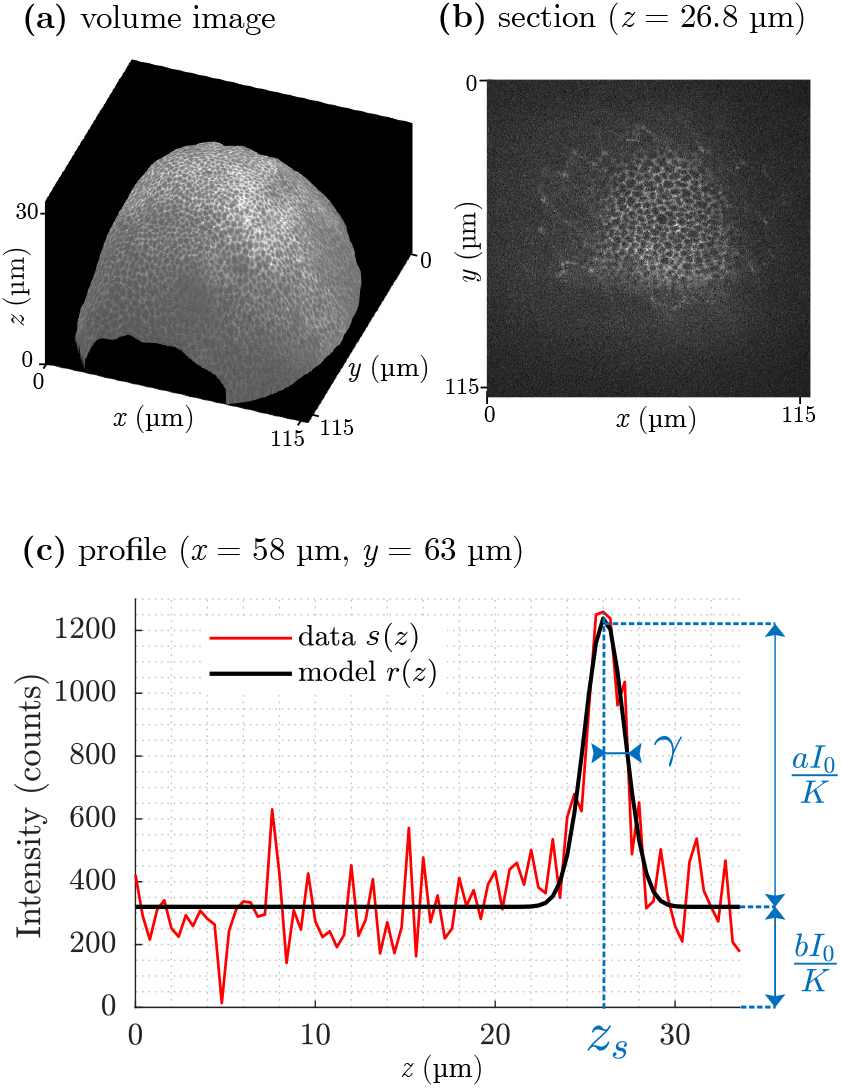
Example of a fluorescent signal coming from an imaginal disk of Drosophila melanogaster embryo. The cell junctions (E-cadherin) have been tagged with GFP to be fluorescent and lie on a thin sheet. The fluorescent signal is acquired using a spinning-disk confocal microscope coupled with an EM-CCD sensor. (a) Volume image of the surface, with (b) a detail of the section at *z* = 26.8 μm, and (c) a profile along *z* (red curve) at (*x, y*) = (58 μm, 63 μm), and the corresponding model of the averaged signal *r*(*z*) (black curve). The position of the peak corresponds to that of the cell sheet at the selected (*x, y*) coordinates. The amplitude of the peak with respect to background is denoted *aI*_0_/*K*, and the background signal is denoted *bI*_0_/*K* where *I*_0_ is the total intensity being sent along *z* axis, and *K* is the number of scanning samples along *z*.

Let us consider the signal *s* along the z-axis (optical axis) at a given lateral coordinates (*x, y*). Figure 1.c (red curve) shows a typical profile of this signal *s* when (*x, y*) belongs to the adherens junction. This signal is sampled over a set of *K* scanning points 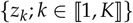, and can be modeled as

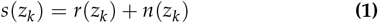

where *r* is a linear combination of a Gaussian shape centered on *z_s_* and of a constant background, as shown in Fig. 1.c (black curve), and *n*(*z_k_*) denotes the noise corrupting the acquisition. More precisely, *r* can be written as

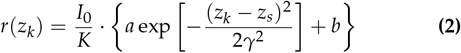

where *I*_0_ is the intensity of excitation light to be distributed along *z*-axis. We assume *I*_0_ is homogeneously distributed so that each point zk received *I*_0_/*K*. The parameter *γ* denotes the standard deviation of the Gaussian shape, and is thus related to the width of the adherens junction. The parameter *b* relates to the proportion of background light reaching the sensor. The parameter *a* is the amplitude of the Gaussian shape with respect to background. Note that the values of *a* can strongly vary with respect to (*x, y*), as for example in Fig. 1.b where only the cell contours contain fluorescent tags.

The signal *r* is corrupted by two sources of noise contained in *n*: i) electronic noise (mainly thermal and readout noise), which can be modeled as white Gaussian noise of constant variance *β,* and ii) signal-dependent noise related to photon noise and amplification of the signal. In this work, we consider that, after amplification, the signal we observe contains a sufficient number of counts, so that this signal-dependent noise can be modeled as a Gaussian noise of variance *αr*(*z_k_*), where *α* is related to the photoelectron amplification. Thus, the *n*(*z_k_*) are assumed to be independent Gaussian random variables, with zero-mean and with variance

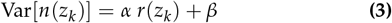

These parameters *α* and *β* allows to cope with various sensor noise models [20–22].

In this paper, the parameters *z_s_, γ, a*, and *b* are the unknown parameters to estimate – although their values might be known to some extent. In the estimation problem, *z_s_* is the parameter of interest, while *γ, a*, and *b* are nuisance parameters. The other parameters constitute experimental parameters.

### B. Cramér-Rao bound

Our goal is to estimate *z_s_* the most accurately possible, having a fixed photon budget *I*_0_ to be distributed on the set of scanning points {*z_k_*; *k* ∈ ⟦1, *K*⟧. Given this photon budget *I*_0_, what are the number *K* and the positions *z_k_* that optimize the estimation accuracy for *z_s_*?

To answer this question, we propose to optimize *K* and the set of *z_k_* with respect to the Cramér-Rao Bound (CRB) of *z_s_*. The CRB represents the lower bound on the estimation variance that can be obtained with an unbiased estimator. As in [23], but considering the noise model given in Eq. (3), the CRB of *z_s_* can be expressed as (see Appendix A)

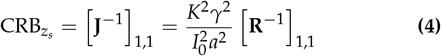

where **J** is the Fisher Information Matrix associated to the estimation of the 4 unknown parameters (*z_s_, γ, a, b*), and where

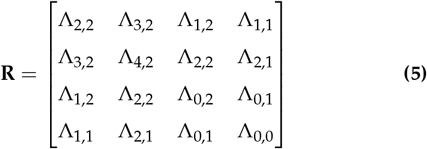

with

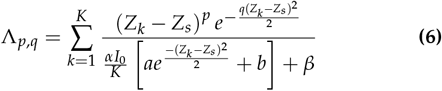

with defining the reduced parameters *Z_s_* = *z_s_*/*γ*, and *Z_k_* = *z_k_/g*.

This expression depends on *z_s_*, which is the parameter of interest, and on the parameters *a, b* and *γ*, which are generally unknown. Note that, usually, an approximate value of *γ* is known, as it corresponds to the measurable thickness of the cell sheet. The expression also depends on the intensity *I*_0_, on the noise parameters *α* and *β*, and on the number K and the positions *z_k_* (*k* ∈ ⟦1, *K*⟧) of the scanning points. Our goal is to analyze how to optimize *K* and *z_k_* to decrease CRB*_z_s__*.

In the following, we will focus on two situations: first, when we have no *a priori* knowledge on the surface’s location *z_s_* (section 3), and second, when we know approximately in which interval *z_s_* has to be searched (section 4).

## 3. OPTIMAL SAMPLING WITHOUT PRIOR ON THE SURFACE LOCATION

In this section, we assume that we have no prior information on the position *z_s_* of the biological surface. In this case, the whole axial range available has to be scanned, with scanning points *z_k_* uniformly distributed along *z*. Our goal is then to find the optimal value of the sampling step, *i.e.* the *spacing* between two consecutive scanning points.

### A. Reformulation of the optimization problem

Let us denote [*z*_min_, *z*_max_] the axial range available. As previously mentioned, the scanning points *z_k_* will be uniformly distributed on [*z*_min_, *z*_max_] so that *z*_*k*+1_ – *z_k_* = *δ*, with *δ* denoting the constant sampling step, and *z*_1_ = *z*_min_ + *δ*/2 and *z_k_* = *z*_max_ – *δ*/2. Let us define *I* = *z*_max_ – *z*_min_ the length of the interval to probe. The number *K* of scanning points is then *K* = *l/δ.* Thus, optimizing the scanning strategy is equivalent to determine the value of *δ* that minimizes CRB*_z_s__*.

Let us define the reduced parameters Δ = *δ/g* and *L* = *l/g*. The coefficients Λ_*p,q*_ of Eq. (6) can be rewritten as

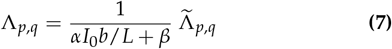

where

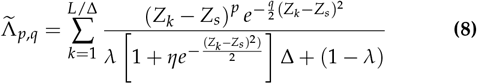

with

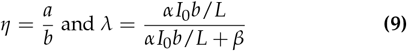

The parameter *η* is the signal to background ratio. The parameter *λ* is a noise coefficient that takes values between 0 and 1. In the expression of *λ*, the quantity *αI*_0_*b/L* corresponds to the variance of the signal-dependent noise that comes from the background, on a typical length of *γ*, and *β* is the variance of the electronic noise. Therefore *λ* can be seen as the proportion of signal-dependent noise with respect to the total amount of noise, considering the background only. When *λ* → 0, *β* » *αI*_0_*b/L*, which means that the electronic noise becomes predominant, whereas, when *λ* → 1, the signal-dependent noise is predominant. It corresponds to the case of an ideal sensor without additive electronic noise. Note that when *λ* = 1, the CRB of *z_s_* corresponds to that obtained in Poisson noise regime.

In the following, we define the matrix 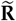 that is of the same form of matrix **R** given in Eq. (5), but filled with the 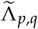 coefficients instead of Λ_*p,q*_. The CRB of *z_s_* becomes

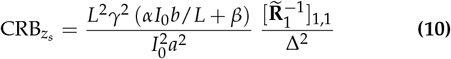

Finding the value of *δ* that minimizes CRB*_z_s__* is thus equivalent to find the value of Δ = *δ/g* that minimizes 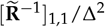.

From Eq. (8), it comes that 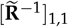 is a function of *Z_s_*, *η, λ*, and of the set {*Z_k_*}_*k*=1… *K*_ i.e. of the set {*Z*_1_, L, Δ}. We can check that 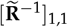 depends no more on *L*, as soon as *L* is sufficiently high (typically *L* = *l/g* ≳ 10). Indeed, increasing *L* while keeping all other parameters constant only improves the estimation of the unknown background parameter *b*, and has no impact on the estimation precision of *a, γ* and *z_s_*.

Because 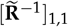 depends on *Z_s_* itself, which is to estimate, we propose to follow a minimax approach. We minimize the maximal CRB – the worst CRB value – over all *Z_s_* values. When ignoring bounding effects obtained when *z_s_* is close to *z*_min_ or *z*_max_ (within a typical distance of 3*γ*), it comes that CRB*_z_s__* varies as a Δ-periodic function of *Z_s_*. We thus only have to consider the maximal value of CRB*_z_s__* over all *Z_s_* in [*Z*_0_ – Δ/2, *Z*_0_ + Δ/2] with *Z*_0_ = (*Z*_max_ + *Z*_min_) /2. We then search for the optimal Δ value

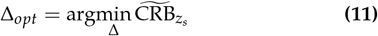

with

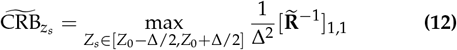

Now, provided *L* is sufficiently high, and ignoring *Z_s_*-bounding effects, Δ_*opt*_ depends only on *λ* and *η.* Through these two reduced parameters *λ* and *η*, Δ_*opt*_ takes into account the parameters of the acquisition and of the model (*i.e. z*_min_, *z*_max_, *I*_0_, *a, b, γ, z_s_, α* and *β*).

### B. Analysis

Figure 2.a shows the evolution of 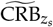 with respect to Δ, when *η* = 1 and for several values of *λ*. Note that when 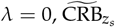 no more depends on *η* (see Eq. (8)). For ease of comparison between different *λ* values, the CRB has been divided by its minimal value 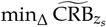.

**Fig. 2.**
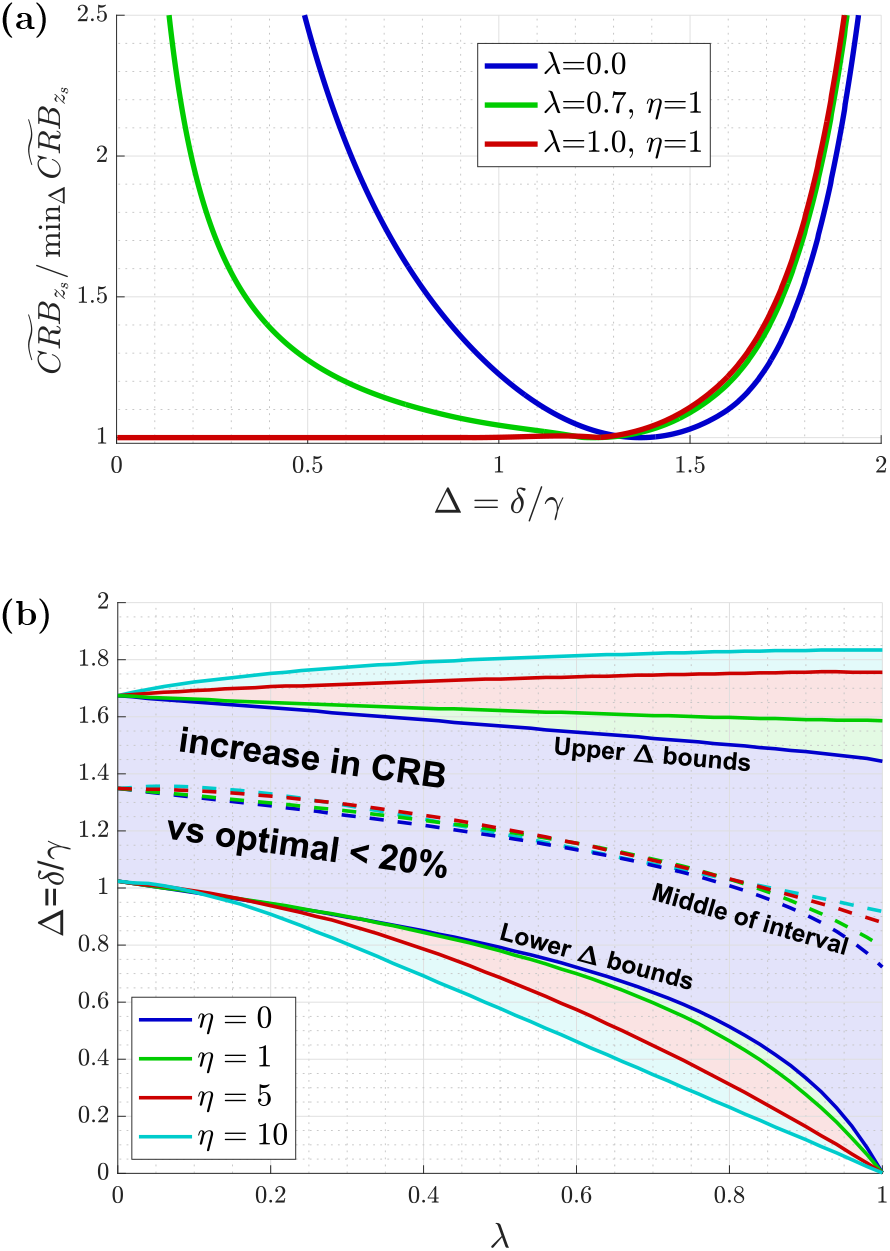
(a) Evolution of 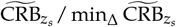 as a function of Δ = *δ/γ* and for several values of *λ* and *η*. (b) Evolution as a function of *λ* (and for several values of *η*) of the range of Δ-values for which the increase in 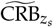 with respect to the optimal value 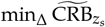 is lower than 20%. Plain curves correspond to the upper and lower bounds of the Δ ranges and dashed curves to the middle of the Δ-ranges.

As can be seen in Fig. 2.a, for all conditions, 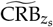 presents a clear minimum, and increases to infinity at large Δ. A contrario, when Δ → 0, the behavior of 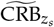 depends on *λ*. When *λ* = 1, CRB*_z_s__* stays almost constant and close to its minimal value, as long as Δ < 1.3 (red curve in Fig. 2.a). When 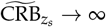 when Δ → 0. Indeed, *λ* < 1 corresponds to *β* > 0, the case of a non-ideal sensor. In this case, when Δ → 0, the number of scanning points increases. Then, because the photon budget is fixed, the number of photons per scanning points tends to zero while the variance *β* of the electronic noise remains constant.

From the position Δ_*opt*_ of this minimum, for each *λ* and *η*, we can derive the optimal sampling step *δ_opt_* = *γ* Δ_*opt*_, provided *γ* is known. Fortunately, in many biological configurations an approximate value of *γ* is generally known, allowing then to derive an approximate value of *δ*_*opt*_. Considering this point, our goal is then not necessarily to determine the precise value Δ_*opt*_ but more interestingly to determine a range of Δ-values, for which the performance are close to the optimal CRB value.

Accordingly, Figure 2.b shows the range of Δ, for which the increase in 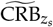 from its minimal value is bounded to 20%, as a function of *λ*. It has been plotted for several values of *η* varying from 0 to 10. Note that although *η* = 0 corresponds to a situation for which it is not possible to estimate *z_s_* (since *a* = 0), it has been plotted as the limit case of null signal-to-background ratio. The larger the area where the increase in CRB is lower than 20%, the more robust to an uncertainty on the parameter *γ*.

Having an approximate value 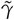 of *γ*, we can set the value of the spacing *δ* so that 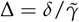 corresponds to the middle of the Δ-range (see dashed lines in Fig. 2.b). Furthermore, although the signal-to-background ratio *η* = *a/b* is another unknown that can strongly varies in the image, it can be seen that the middle of this Δ-range is almost independent of *η*, even when *η* is varying from 0 to 10. Furthermore, it can be noted that setting for example Δ = 1.3 is a robust solution, whatever the value of *λ*.

Let us now analyze the robustness of this approach to an error on *γ*. We denote *γ* the true standard deviation of the Gaussian, and 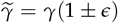 the erroneous value used to define the scanning scheme. With these notations, the reduced parameter Δ = *δ/γ* is changed into 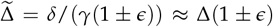. This means that Δ has the same relative error *ϵ* as *γ*. Figure 3 shows how an error on *γ* affects the estimation of *z_s_* depending on noise parameters. To do so, we first set Δ near its optimal value, i.e. in the middle of the Δ-range aforementioned (where the increase in CRB with respect to optimal is lower than 20% - see Fig. 2.b). We then plot the evolution of 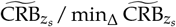 as a function of the noise parameter *λ*, and computed for 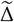. The curves are plotted for several values of the relative error on *δ* (*ϵ* = 0%, 10%, 20%, 30%) and for several values of *η*. Without any error on *γ* (blue curves), choosing Δ in the middle of the 20% interval leads to a very limited increase in CRB from optimal. With a large 30% error on *γ* (yellow curve), the increase in CRB with respect to optimal remains bounded by a factor of 1.4 (1.15 for an error of 20%, see red curve). When *λ* → 1, i.e. as the contribution of the electronic noise *β* decreases, the CRB ratio tends to 1, which confirms the robustness of this approach to a certain mis-knowledge on *γ*.

**Fig. 3.**
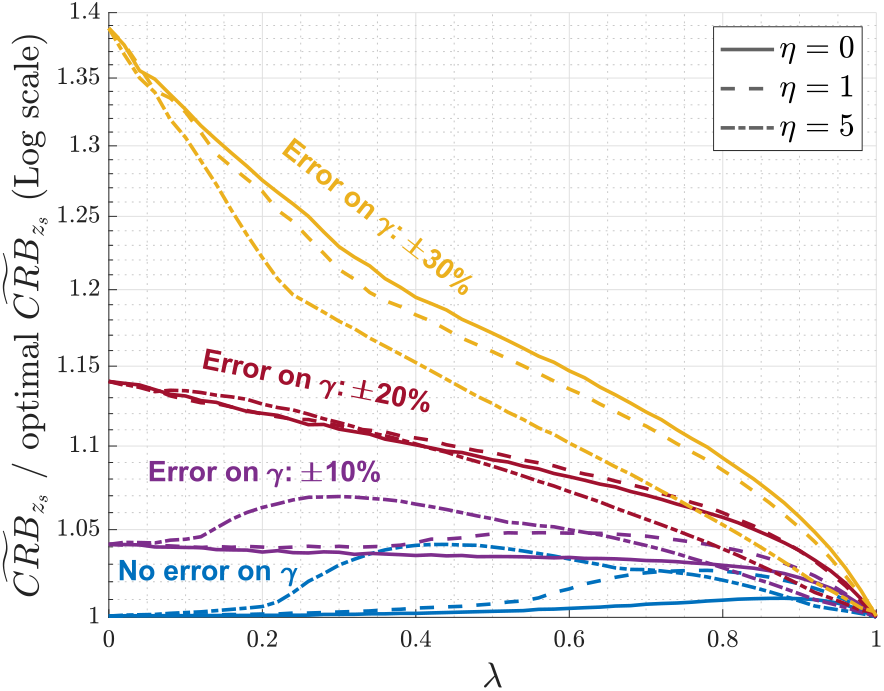
Robustness to an error on *γ*. Evolution of 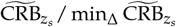, as a function of *λ* in the ideal case where *γ* is perfectly known (in blue) and in case of an error on *γ* of 10% (purple), 20% (red) and 30% (yellow). These curves have been plotted for *η* = 0 (plain lines), *η* = 1 (dashed lines) and *η* = 5 (dash-dotted lines).

In situations in which the *a priori* knowledge on *γ* is not accurate enough, estimation performance may be far from optimal. To compensate this decrease of performance with respect to optimal, a possibility is to increase *I*_0_. However, it results in increasing phototoxicity as well. In such situations, it can be interesting to use an iterative procedure. Rather than directly sending the whole photon budget *I*_0_ on the tissue, we can perform a first acquisition with lower intensity, and with *δ* determined from Fig. 2.b using the user’s prior on *γ*. This will then allow to perform further acquisitions with more accurate priors on *γ*, and thus with values of 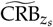 closer to optimal. Last but not least, such iterative approaches will also progressively provide a prior knowledge on the location *z_s_*. This latter point has thus to be taken into account in the design of the scanning strategy, and will be analyzed in the next section.

## 4. OPTIMAL SAMPLING WITH PRIOR ON THE SURFACE LOCATION

In the previous section, we assumed no prior information on the position *z_s_*. This led to a uniform distribution of the scanning points over the whole *z*-range. On practical terms however, one can sometimes have an approximate prior knowledge of *z_s_*. This prior may come from previous estimation at neighboring points (*x, y*) or in the context of an iterative procedure (at given (*x, y*), an approximate estimation of the surface’s location *z_s_* precedes a more accurate one). When *z_s_* is approximately known, instead of scanning all the available *z*-space, we rather perform a few acquisitions around the approximate value of *z_s_*.

In this section, we still assume that the *K* scanning points are regularly spaced by a constant *δ*. We study the influence of *K* and *δ* on the estimation performance, knowing that the total span of the scanning points no more needs to cover the full z-range [*z*_min_, *z*_max_].

### A. CRB expression

The scanning points *z_k_* are positioned around the a priori value of *z_s_*. We denote z0 the center of the acquisition interval [*z*_1_; *z_k_*], defined as *z*_0_ = (*z_k_* + *z*_1_)/2. Our goal is to determine the optimal values of *K, δ*, and of the position *z*_0_ of the interval [*z*_1_, *z_k_*] with respect to the amount of prior knowledge we have on *z_s_*.

In this case, the CRB of *z_s_* can be written as

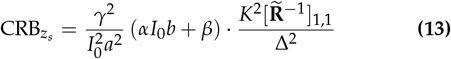

where the coefficients 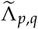 of the matrix 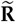 take a different form than that of in the previous section:

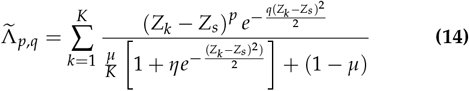

with

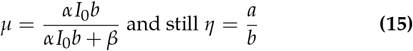

We have introduced a new parameter *μ* that replaces the parameter *λ* of section 3. Indeed, *λ* depends on the span *Kδ* that is no more fixed to *l* = *z*_max_ – *z*_min_. The parameter *μ* now represents the proportion of the signal-dependent noise with respect to the total amount of noise, considering the background signal only.

Thus, the quantity CRB_*z_s_*_ depends on the scanning strategy (*K, δ*, and *z*_0_) only through the quantity 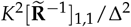, which depends on the reduced parameters *μ, η, Z_s_* and of course on *Z*_1_, *Z*_2_,…, *Z_k_* (i.e. on Δ = *δ/γ, K*, and *Z*_0_ = *z*_0_/*γ* the middle of [*Z*_1_; *Z_K_*]). In the following, the goal is to search for the optimal values of Δ, *K* and *Z*_0_ that minimize this CRB.

The Fisher Information Matrix is non-invertible when *K* < 3 and in some situations when *K* = 4. We thus start our analysis with *K* = 5, as it corresponds to the simplest situation.

### B. Analysis of the CRB when *K* = 5

Let us assume first that *z_s_* is perfectly known. Figure 4.a shows the evolution of CRB_*z_s_*_ / min_Δ,*z*_0__ CRB*_z_s__*, for *K* = 5 as a function of *Z_s_* – *Z*_0_ and of Δ. This normalized CRB depends on *μ, η* and on the acquisition parameters through the reduced variables *Z_s_* – *Z*_0_ = (*z_s_* – *z*_0_)/*γ* and Δ = *δ/γ*. The black-contour plots correspond to *μ* = 0 (electronic noise only) and to a signal-to-background ratio of *η* → 0. The red-contour plots correspond to *μ* = 1 (pure signal-dependent noise) and *η* → ∞. Whatever the values of *η* or *μ*, the minimum of CRB*_z_s__*, is reached for *Z*_0_ = *Z_s_*. Then, we have min_Δ,*Z*_0__ CRB*_z_s__*, = min_Δ_ CRB*_z_s__,|*z*_0_=*z*_0_*, and Δ_*opt*_ = argmin_Δ_ CRB*_z_s__*, |*z*_0_=*Z_s_*.

**Fig. 4.**
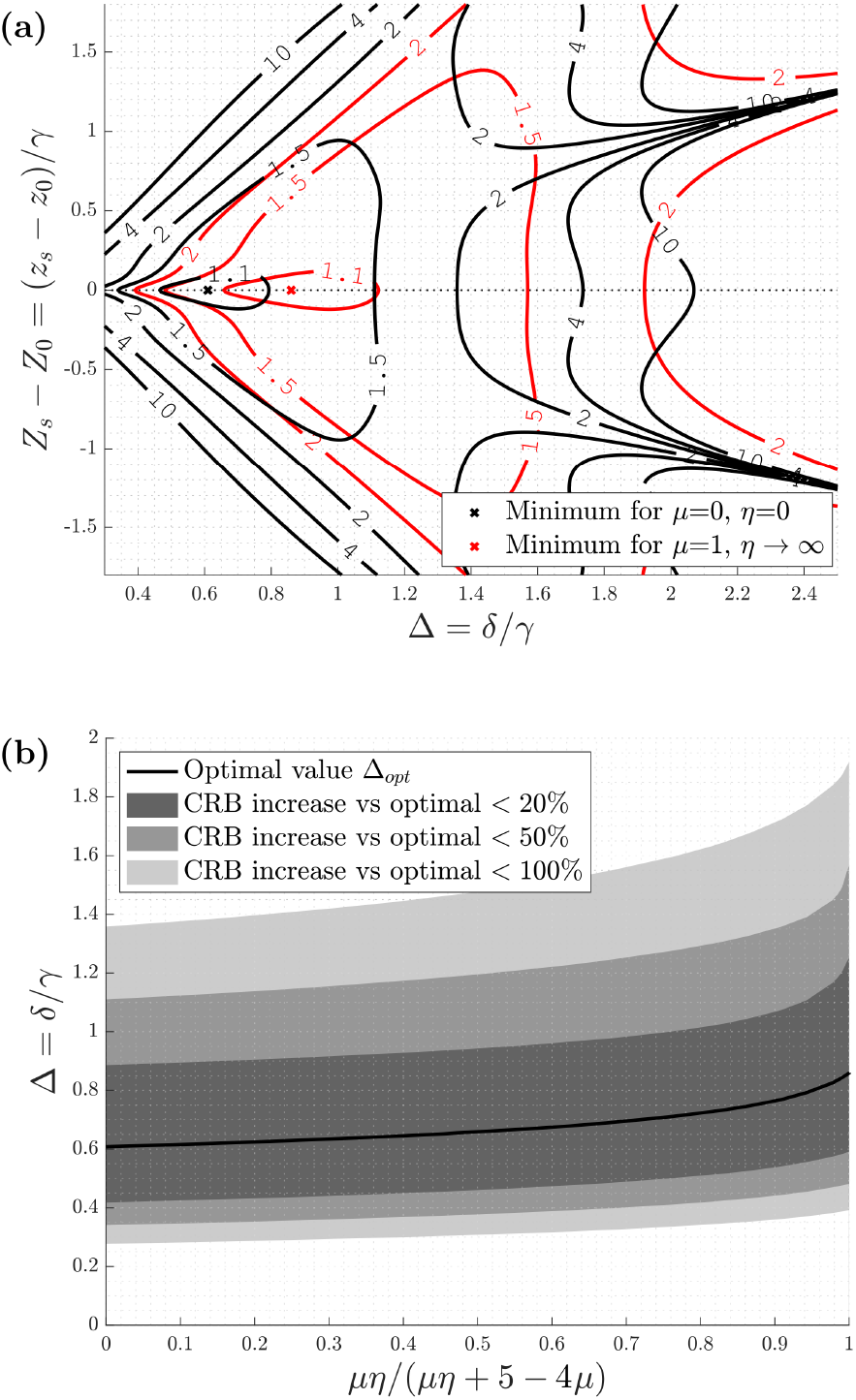
Analysis when only *K* = 5 scanning points are used: (a) CRB*_z_s__* / min_Δ,*z*_0__ CRB*_z_s__* as a function of Δ and of the reduced variable *Z_s_* – *Z*_0_, with *Z*_s_ = *z_s_/γ* and *Z*_0_ = *z*_0_/*γ*, *z*_0_ being the center of the sampling interval. The black-contour plots correspond to the case where *μ* = 0 and *η* → 0, while the red-contour plots correspond to the case where *μ* = 1 and *η* → ∞. The two crosses correspond to the locations where the CRB for both cases take their minimum. (b) When *z*_0_ = *z_s_*, evolution with respect to the reduced variable *μη*/ (*μη* + 5 – 4*μ*) of the optimal value Δ_*opt*_ and of the Δ-ranges for which the CRB increase is lower than 20%, 50 % and 100 % of its minimal value.

Moreover, when *K* = 5, we can write from Eq. (13) and Eq. (14) the quantity minA CRB*_z_s__*, |*z*_0_=*Z_s_* as a function of a single reduced coefficient *μη*/(*μη* + 5 – 4*μ*) instead of both *μ* and *η*. Figure 4.b shows the optimal spacing Δ_*opt*_ as a function of this reduced parameter when *K* = 5 and *Z*_0_ = *Z_s_* (black curve). We have also plotted the range of Δ values for which the increase in CRB remains within 20%, 50%, and 100% from the minimal CRB. It appears from these results that in situations where *μ* or *η* are not exactly known, setting for example Δ = *δ/γ* = 0.7 can be a robust solution, for which the increase in CRB always stays lower than 20%.

In Fig. 4.b, we set *Z*_0_ = *Z_s_*. It is of course not realistic, since *z_s_* is the parameter to estimate. Since *z_s_* – and thus *Z_s_* – is only approximately known, we introduce 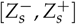, which is the interval of possible *Z_s_*-values. In order to take this interval into account in the optimization of the acquisition parameters, we still follow a minimax approach: we minimize the worst CRB value over all *Z_s_* taking value in 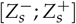. In other words, the optimization problem consists in finding the acquisition parameters Δ, *K, Z*_0_) that minimize

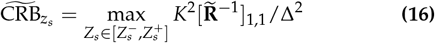

As the CRB is symmetric with respect to *Z*_0_ – *Z_s_* (see Fig. 4.a), *Z*_0_ has to be set in the middle of 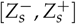. In the following, we will thus fix *Z*_0_ to 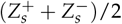.

In Fig. 5, we have plotted the evolution of 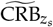 as a function of the uncertainty 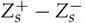, *i.e.* the length of 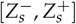(see blue curve for *K* = 5), when the signal-to-background ratio is set to *η* = 2 and the noise parameter to *μ* = 0.7. This CRB has been computed for Δ = 0.7, which is close to the optimal value Δ_opt_ when *Z*_0_ ≈ *Z_s_* and *K* = 5 (see Fig. 4.b). The signal-to-background ratio is set to *η* = 2, and the noise parameter is *μ* = 0.7. It can be seen that the CRB increases quickly as the uncertainty 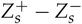 increases.

**Fig. 5.**
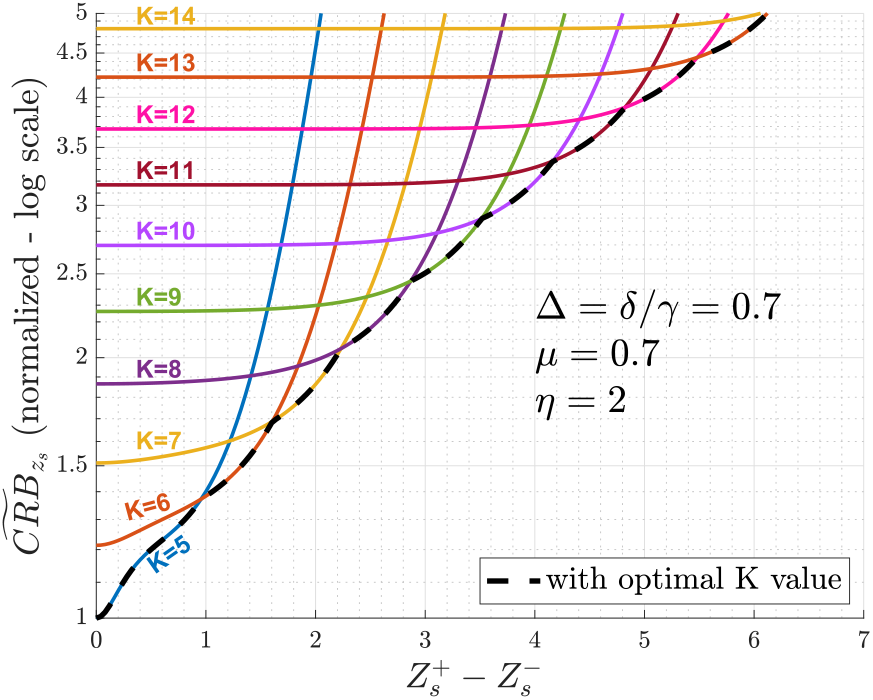
Example, for *μ* = 0.7 and *η* = 2, of the evolution of 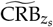 as a function of the uncertainty on *Z_s_* (i.e. the value 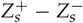), when Δ is fixed to 0.7, and when *K* progressively increases from *K* = 5 to *K* = 14. This CRB has been normalized so that its minimum value (reached when 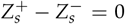 and *K* = 5) is equal to 1. The value of 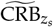 computed with optimal *K* value has been super-imposed in black dashed line.

### C. Analysis of the CRB in the general case

A solution to limit the increase in CRB is to add new scanning points while keeping Δ constant. Netherless, it results in the decrease in the intensity sent per each scanning point, as one operates at constant photon budget.

In Fig. 5, the evolution of 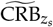 obtained for *K* = 6,7,…, 14 has then been added to the curve obtained for *K* = 5. The lower envelop of all curves (dashed line) corresponds to 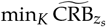, i.e. to the values of 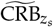 for which the best K-value has been computed for each 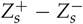. We can see that changing the number *K* of scanning points allows to limit the increase in 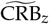 when the uncertainty on *Z_s_* increases.

So far, in Fig. 5, we have fixed Δ to 0.7. We must address whether this choice on Δ is still relevant when the uncertainty on *Z_s_* increases. We have thus plotted in Fig. 6.a the evolution of 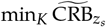 (i.e. of 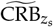 with *K* set to its optimal value) for several Δ values, and still when *μ* = 0.7 and *η* = 2. The lower envelope (red curve) provides an overall optimal CRB, for which the best *K* and Δ are selected for each uncertainty value 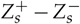. This envelope corresponds to min_Δ,*K*_ 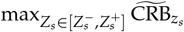 (with Δ = *δ/γ* varying between 0.01 and 2 with a step of 0.01 for this curve). It appears that the optimal couple (*K*, Δ) changes rapidly with the uncertainty 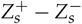. Thus, one cannot rely on a single Δ to be optimal in a wide range of *Z_s_* uncertainty. One rigorous but fastidious way to optimize the estimation would be to provide the best couple of parameters Δ and *K* for each 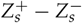, and of course for several values of the parameters *μ* and *η*.

**Fig. 6.**
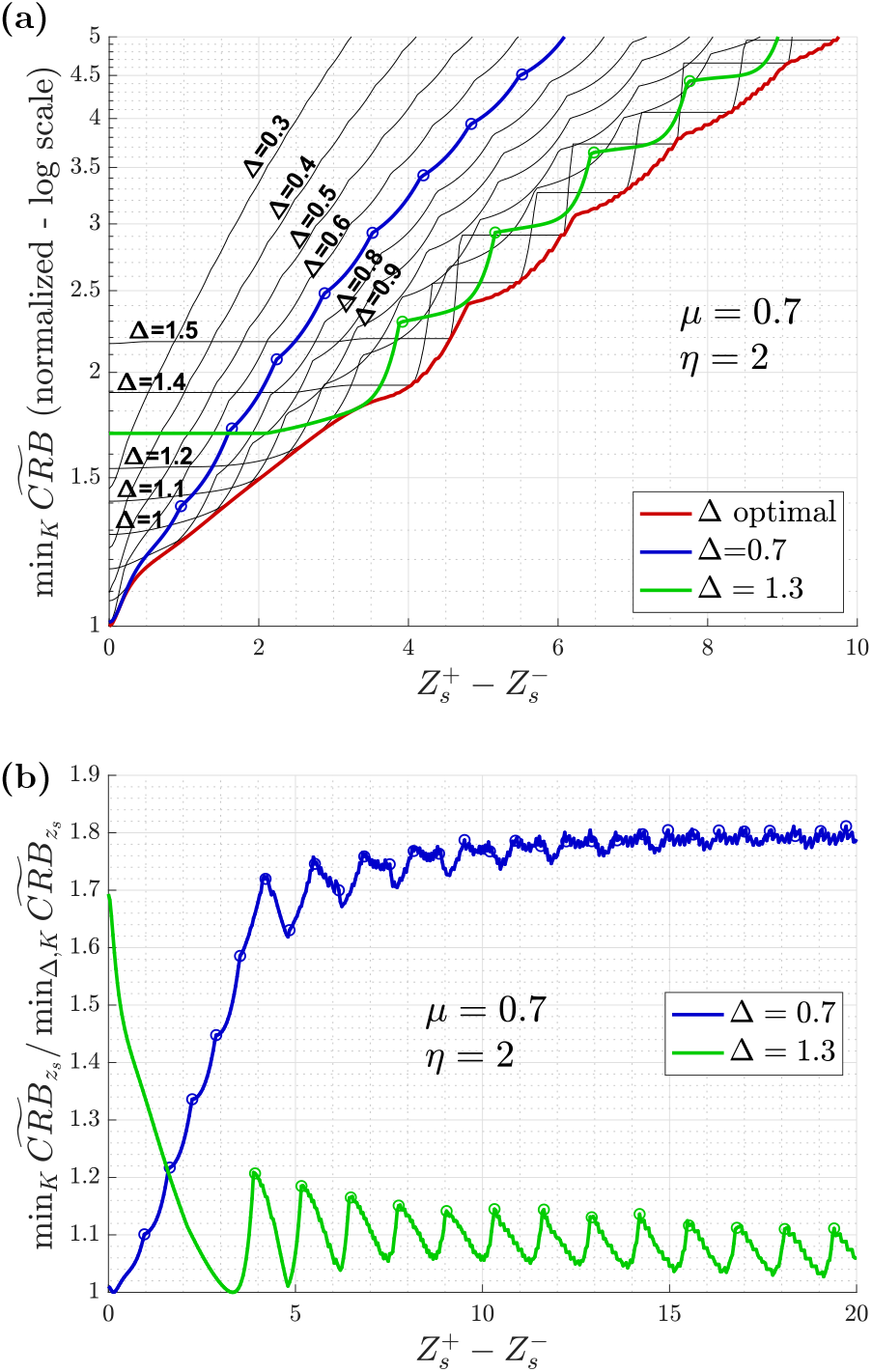
Example for *μ* = 0.7 and *η* = 2: (a) evolution of 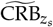 as a function of the uncertainty on *Z_s_*, for different values of Δ and with selecting for each uncertainty value the optimal number *K* of scanning points. This CRB has been normalized so that its minimum value (reached when 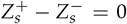 and *K* = 5) is equal to 1. The blue line corresponds to Δ = 0.7 and the green one to Δ = 1.3. The circle points indicate where the optimal values of *K* increase by 1. The red line corresponds to the optimal setting of both *K* and Δ among all possible values (with Δ sampled with a step of 0.01). (b) Evolution of minK 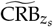 divided by the optimal CRB value 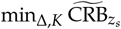 (optimized both on *K* and Δ), as a function of the uncertainty 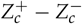 on *Z_s_*, when Δ is fixed to 0.7 (blue curve) and to 1.3 (green curve).

To further gain insight in the choice of Δ, we emphasize the case Δ = 0.7 (in blue on Fig. 5.b), which has been shown to be quasi-optimal when *Z*_0_ ≈ *Z_s_* and *K* = 5, whatever the values of *μ* and *η* (see Fig. 4.b). As expected, the curve appears near optimal at low values of 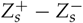, but increases much faster than optimal for 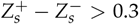. As 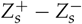 becomes larger, the number of scanning points has to be increased, and we gradually approach the case of estimating *z_s_* without prior, dealt in the previous section. A large number of scanning points at constant photon budget implies fewer photons per points. The noise variance is then increasingly dominated by the variance *β* of the electronic noise. Therefore, when 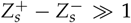, the optimization becomes equivalent to the case without prior and *λ* → 0. For this case, the optimal choice was around Δ = 1.3 (see previous section). The curve for which Δ = 1.3 has thus been underlined in Fig. 6.a (in green).

These two extreme settings Δ = 0.7 and Δ = 1.3 lead to performance close to those obtained with optimal Δ and *K* settings (red curve in Fig. 6.a). In Fig. 6.b, we have plotted the ratio of 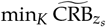 (i.e. 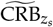 with optimized *K* value) for Δ = 0.7 or Δ = 1.3, over 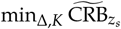 (i.e.. 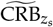 with both optimal Δ and *K* settings), still when *μ* = 0.7 and *η* = 2. It appears that if we choose Δ = 0.7 when 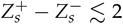 and Δ = 1.3 otherwise, the cost with respect to the optimal 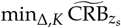 value remains lower than a factor 1.21 i.e. the increase in CRB from optimal will stay under 21%.

### D. Simplified two-spacing strategy

Following this analysis, we propose an alternative strategy to the systematic determination of the optimal parameters Δ and *K*. This simpler approach is to only consider the setting Δ = 0.7 for small values of the uncertainty 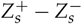, and Δ = 1.3 otherwise. There remains now to determine the value 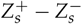 where the transition between these two solutions occurs. We have seen that this transition happened for 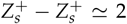 when *μ* = 0.7 and *η* = 2 (see Fig. 6.b). In Fig. 7.a, we have plotted this transition with respect to *μ*, and for both *η* = 5 and *η* → 0. For values of 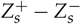 below this transition, the scanning has to be done with Δ = 0.7, while it has to be performed with Δ = 1.3 above this transition. Furthermore, the optimal values of *K* corresponding to each situation have also been reported on this graph.

**Fig. 7.**
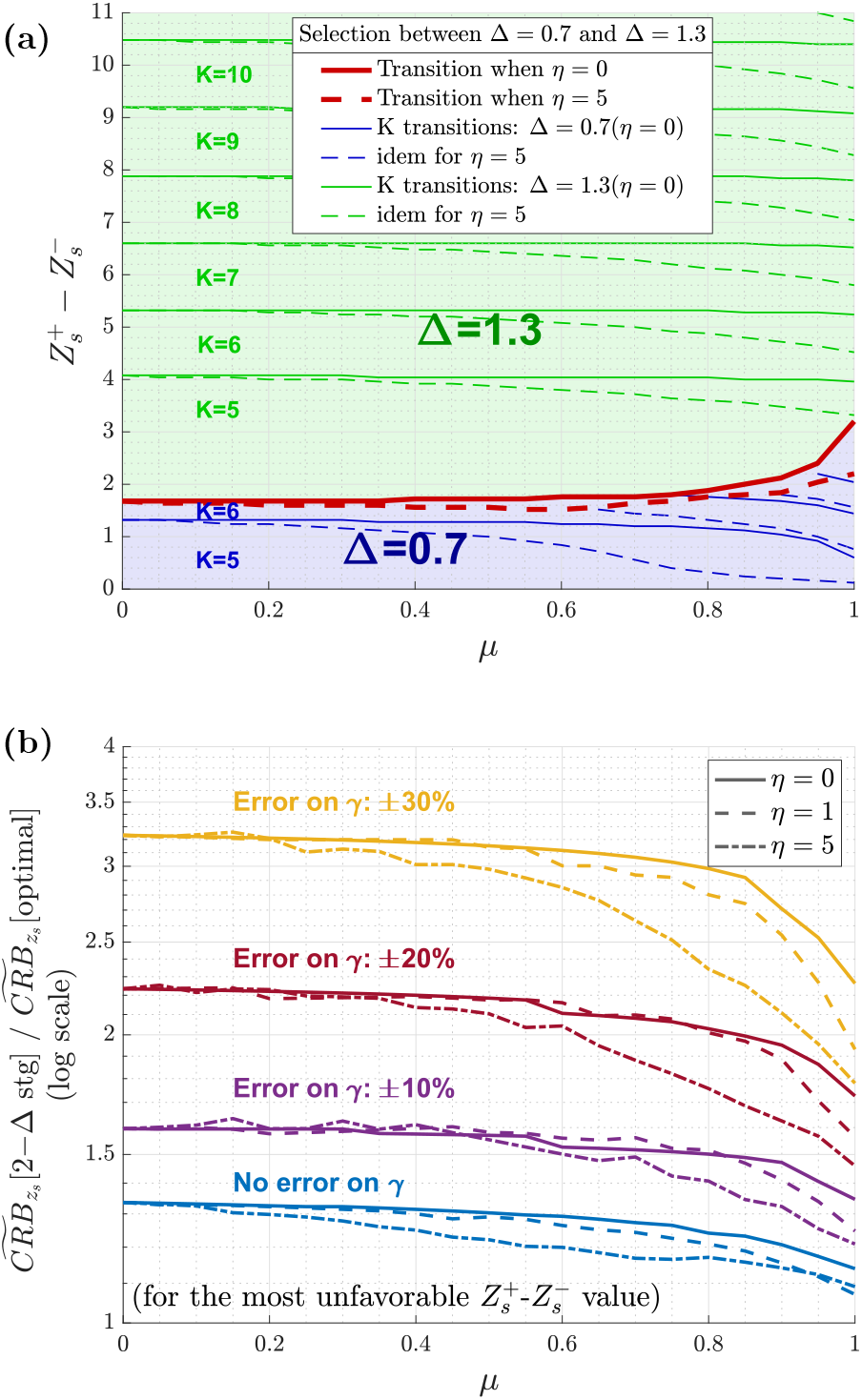
(a) Selection between A = 0.7 (blue zone) and Δ = 1.3 (green zone) as a function of *μ* and of the uncertainty on *Z_s_* (i.e. 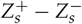). The frontier between these two situations has been plotted with a red line. The optimal choice of *K* has also been reported on this graph (with frontiers marked with green or blue lines). These curves have been plotted for *η* = 0 (plain lines) and *η* = 5 (dashed lines). (b) Robustness to the knowledge of *γ*. Evolution, as a function of *μ*, of the ratio of 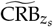 obtained using the approach proposed in (a) to that of obtained with optimal Δ and *K* values. The ratio is computed for the most unfavorable uncertainty 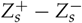, i.e. the uncertainty that leads to the highest CRB ratio. This ratio has been plotted first in the ideal case where *γ* is perfectly known (in blue), and then in the case of an error on *γ* of 10% (purple), 20% (red) and 30% (yellow). These curves have been plotted for *η* = 0 (plain lines), *η* = 1 (dashed lines) and *η* = 5 (dash-dotted lines).

Considering this simplified strategy, a question arises: how much do we lose by using this simplified strategy instead of using the strategy with optimal Δ and *K*? To address this point, we show in Fig. 7.b (blue curves) the ratio of 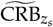 obtained with the simplified strategy and of 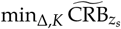 obtained with optimal Δ and *K* settings. This ratio depends on *η, μ* and 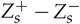. For the sake of simplicity, we have plotted the worst CRB ratio i.e. the highest ratio over all values of 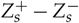 as a function of the noise parameter *μ*, and for several *η* values. It appears that the increase in CRB is very limited. The maximal increase in CRB is obtained for *μ* = 0, with a ratio of 1.32 (see blue curve in Fig. 7.b). Note that, to keep performance level close to optimal, this can be compensated by increasing the photon budget *I*_0_ by 32%.

Finally, we investigate the robustness of this simplified strategy to an error on *γ*. In Fig. 7.b, we have added the ratio when assuming an error *ϵ* on *γ* of 10% (violet curves), 20% (red curves) and 30% (yellow curves). It can be seen that the consequences of a mis-knowledge of *γ* in the case of a prior are more important than that of without prior (cf. previous section and Fig. 3). For example, an error on *γ* of 30% leads this time to an increase in the CRB by a factor of 3.2, which is to compare with an increase by a factor of 1.4 when having no prior on *z*_s_. This difference comes from the fact that this time, an error on *γ* will result in an error on the transition point between the Δ = 1.3 and Δ = 0.7 domains, but also on the determination of K. Nevertheless, it should be pointed out that as soon as a prior on *z_s_* is available, a quite good prior on *γ* can be expected thus limiting the error on *γ* and the increase in CRB.

## 5. CONCLUSION

In this paper, we have proposed a CRB-based approach to optimize the scanning strategy along z-axis, given a fixed photon budget, when the goal is to determine the location *z_s_* of a labeled biological surface. Here, the fluorescent signal was modeled as a Gaussian shape centered on *z_s_* and corrupted by additive signal-dependent noise. The problem of estimating *z_s_* at fixed photon budget, depends on numerous parameters: parameters related to the acquisition process (noise, photon budget, scanning strategy) and parameters related to the model, including *z_s_* itself. It also depends on the amount of prior knowledge we have on *z_s_*. We have shown that, by introducing a controlled loss in the estimation accuracy, we can design general scanning strategies that only depend on a few reduced parameters.

Without prior knowledge on *z_s_*, the strategy is to scan regularly the full *z*-space. In this situation, the optimal spacing *δ* between two consecutive scanning points only depends on *γ*, which is related to the width of the tissue modeled by the Gaussian shape, on the signal-to-background ration *η*, and on the noise parameter *λ*, which account for the proportion of pure signal-dependent noise with respect to the total amount of noise in the acquisition process. Moreover, when the spacing is set to 1.3*γ*, the increase in CRB from its optimal value will always stay under 20%, whatever the value of the parameters *λ* and *η*. When prior knowledge on *z_s_* is available, a quasi optimal strategy has been proposed in which two situations may occur. If the interval of a priori *z*_s_-values is thinner than approximately 2*γ*, the scanning point should be separated by 0.7*γ*. Otherwise, they should once again be separated by 1.3*γ*.

As a perspective, future work will need to implement and test experimentally the proposed scanning strategy on real biological tissues. The main advantage of this strategy is that it can be easily implemented in 3D-scanning microscopes with targeted illumination such as proposed in [12, 24, 25]. Moreover, an adaptive scanning scheme can be derived from the proposed approach. An estimation of *z_s_* with its associated CRB can be converted to a prior for the next round of estimation. The location of the surface can thus be recovered for each selected coordinates (*x, y*). Furthermore, the precise choice of (*x, y*) that achieves the best trade-offs between accuracy and photon budget is an interesting perspective of this work. Eventually, it is worth noting that this paper focused on the estimation accuracy of *z_s_* only, even though the whole parameters of the model – including the width of the tissue – were estimated simultaneously. It would be interesting to draw scanning schemes that consider jointly the estimation accuracy of the position and of the width of the biological surface.

### A. FISHER INFORMATION MATRIX FOR *Z_s_* ESTIMATION

Let ***δ*** = (*z_s_, γ, a, b*)^*T*^ the model parameter vector to estimate. The log-likelihood of the acquired sample 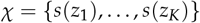 is equal to

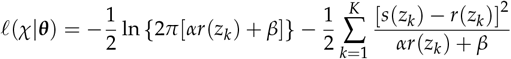

The Fisher information matrix **J** is the 4 x 4 matrix so that, for *n, m* ∈ ⟦1, 4⟧,

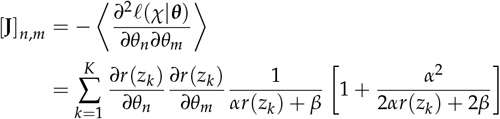

with *δ_n_* denoting the *n*^th^ component of vector ***δ***. As in [15], as soon as the number of counts on the sensor is not too small, the Fisher information matrix **J** becomes

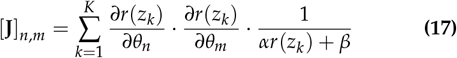

with

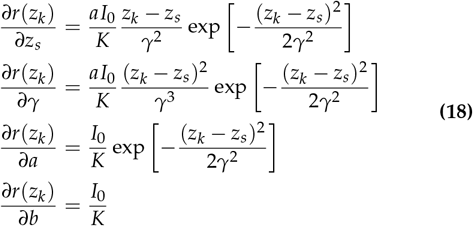

Following Eq. (17) and Eq. (18), **J** can be written as

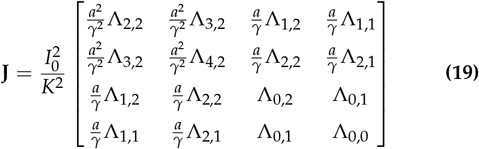

with λ_*p,q*_ expressions given in Eq. (6). The CRB on *z_s_* is thus

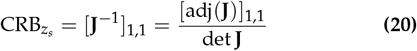

where adj(**J**) denotes the adjugate matrix of **J**, and det(**J**) denotes the determinant of **J**. The cofactor [adj(**J**)]_1,1_ is the determinant of the submatrix of **J** formed by removing the first row and first column. By developing the expression of the determinants of both matrices, and by using the expression of **R** given in Eq. (5), we obtain that 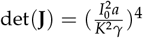 and 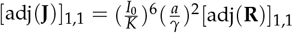, which lead to Eq. (4).

## Funding

Agence Nationale de la Recherche (ANR-18-CE13-028, ANR-17-CE30-0007); Excellence Initiative of Aix-Marseille University - A*Midex (capostromex), a French Investissements d’Avenir programme; The project leading to this publication has received funding from the “Investissements d’Avenir” French Government program managed by the French National Research Agency (ANR-16-CONV-0001, ANR-21-ESRE-0002) and from Excellence Initiative of Aix-Marseille University - A*MIDEX.

## Acknowledgments

We thank Philippe Réfrégier, Sophie Brasselet, Hervé Rigneault and Vincent Bertrand for fruitful discussions on the project.

## Disclosures

The authors declare no conflicts of interest.

## Data availability

Data underlying the results presented in this paper are not publicly available at this time but may be obtained from the authors upon reasonable request.

## Notes

### Competing Interest Statement

The authors have declared no competing interest.

